# Talin-1 determines the direction of primary mouse neutrophils migrating in vivo

**DOI:** 10.64898/2026.05.27.727970

**Authors:** Yan Wang, Qingkang Lyu, Smriti Parashar, Mikhail Fomin, Pei Xiong Liew, Mark H. Ginsberg, Klaus Ley

**Affiliations:** Immunology Center of Georgia (IMMCG), Augusta University, Augusta, GA, USA; Department of Medicine, University of California, La Jolla, CA 92093, USA

## Abstract

Talin-1 is essential for β2 integrin activation in neutrophils, yet its dynamic behavior during neutrophil trafficking in vivo remains poorly understood. Here, we generated EGFP-talin1 knock-in mice, enabling real-time visualization of talin-1 dynamics under physiological conditions. EGFP-talin1 is robustly expressed and preserves without altering β2 integrin expression and activation. Using total internal reflection fluorescence (TIRF) microscopy under flow, we found that talin-1 was rapidly recruited to the plasma membrane during rolling and accumulates further during neutrophil arrest. Intravital microscopy revealed highly dynamic and stage-specific talin-1 redistribution during luminal crawling, transendothelial migration, and interstitial migration. Talin-1 preferentially accumulated at endothelial contact sites during crawling and polarized toward the leading edge during directional migration. These findings establish EGFP-talin1 knock-in mice as platform for visualizing integrin-associated cytoskeletal dynamics in vivo and identify dynamic talin-1 polarization as a feature of neutrophil trafficking.

## Introduction

Talin-1 is a key integrin adaptor protein that links the cytoskeleton to the plasma membrane and to integrins. Talin-1 is required for early embryo development, as constitutive talin-1 knockout mouse embryos die during gastrulation. ^1,2^. Talin’s intracellular localization has been studied in established adherent cell lines, such as mouse embryonic fibroblasts and NIH3T3 fibroblasts, and has been shown to dynamically recruit to nascent adhesion at the cell-substrate interface and to the leading edge during in vitro migration ^3,4^. However, the spatiotemporal distribution of talin-1 in live primary neutrophils under physiological in vivo conditions remains entirely uncharacterized.

Talin-1 is a large cytoskeletal protein encoded by the *Tln1* gene on mouse chromosome 9. The transcript spans 8,559 nucleotides across 57 exons and encodes a 2,541-amino acid protein. Structurally, talin-1 is composed of an N-terminal head domain (∼ 50 kDa), which contains a FERM domain subdivided into F0, F1, F2, and F3 subdomains ^5^, and a C-terminal rod domain (∼ 220 kDa) comprising 13 α-helical bundles (R1-R13). The rod domain includes a linear C-terminal region with two actin binding sites (ABS2 and ABS3) and a C-terminal dimerization site ^6,7^. The F3 subdomain of the FERM head directly binds a conserved NPXY motif in the integrin β cytoplasmic tail of most integrins including β2 integrins ^8,9^. The F3 fold structurally resembles a phosphotyrosine-binding (PTB) domain, allowing high-affinity recognition of the β-tail motif. Crystallographic ^10^ and cryoEM structures ^11,12^ suggest that the talin-1 head domain adopts a linear conformation that lies flat along the inner leaflet of the plasma membrane when bound to β2.

Integrin activation is a tightly regulated process involving both inside-out and outside-in signaling. Partial β2 integrin activation is physiologically initiated during rolling, where engagement of P-selectin glycoprotein-1 triggers a signaling cascade ^13,14^ that results in extension of the integrin extracellular domains but does not induce the high affinity conformation. Functionally, this results in β2 integrin-dependent slow rolling in vivo and in vitro ^15,16^. β2 integrin extension results in conformational changes that expose neoepitopes. Extension of human β2 integrin expose a binding site for the conformation-specific monoclonal antibody KIM127 that recognizes an epitope in the β2 integrin knee region that becomes accessible upon extension (E+ conformation) ^17^.

Rolling human neutrophils can bind to KIM127, which directly demonstrates that rolling is sufficient to induce extension ^13^. However, rolling neutrophils cannot bind mAb24, which binds an epitope in the β2 I-like domain that is exposed when the internal ligand in the αI domain binds to the β I-like domain, indicating the high-affinity (H+) conformation ^18–20^. The H+ conformation can be induced by chemokines and other chemoattractants ^21,22^ and by outside-in signaling ^23^. Only E+H+ β2 integrins can bind ligands in cis, for example, ICAM-1 on endothelial cells ^24^. mAb24 and KIM127 do not block β2-mediated adhesion and do not interfere with each other’s binding or with ligand engagement ^25^. Since neither KIM127 nor mAb24 can bind mouse β2 integrins, we recently generated human β2 integrin (hITGB2) knock-in mice ^26^. Using these mice, β2 integrin conformation can be monitored in real time via live cell imaging.

The present study was designed to track endogenous talin-1 dynamics in primary mouse neutrophils in vivo. To achieve this, EGFP was inserted into exon 4 of the *Tln1* gene (Fig. 1 A). EGFP is linked to the F0 domain of full-length talin-1 via a flexible GGGGS linker. We focused on neutrophils due to their well-studied and tightly regulated integrin activation during recruitment ^27–29^. In circulation, neutrophils remain largely non-adhesive, with minimal interaction with each other ^30^ or other blood cells ^31,32^. At sites of inflammation, they engage in selectin-dependent rolling along the endothelial wall of postcapillary venules ^33,34^. During this phase, transient interactions between partially activated β2 integrins and endothelial ligands slow rolling velocity ^13,35^. Because talin-1 is known to be required for slow rolling ^35^, we hypothesized that talin-1 may be recruited to the plasma membrane during this slow rolling phase and may contribute to the transition to firm arrest. Arrest from rolling is typically triggered by chemokines that bind G-protein-coupled receptors on rolling neutrophils, leading to rapid and full activation of β2 integrins, inducing the extended high-affinity (E+H+) conformation ^27,36^. It is known that talin-1 is involved in the transition to the E+H+ integrin conformation ^35^, but it is not known where it is located in the arresting cell.

**Figure 1.**
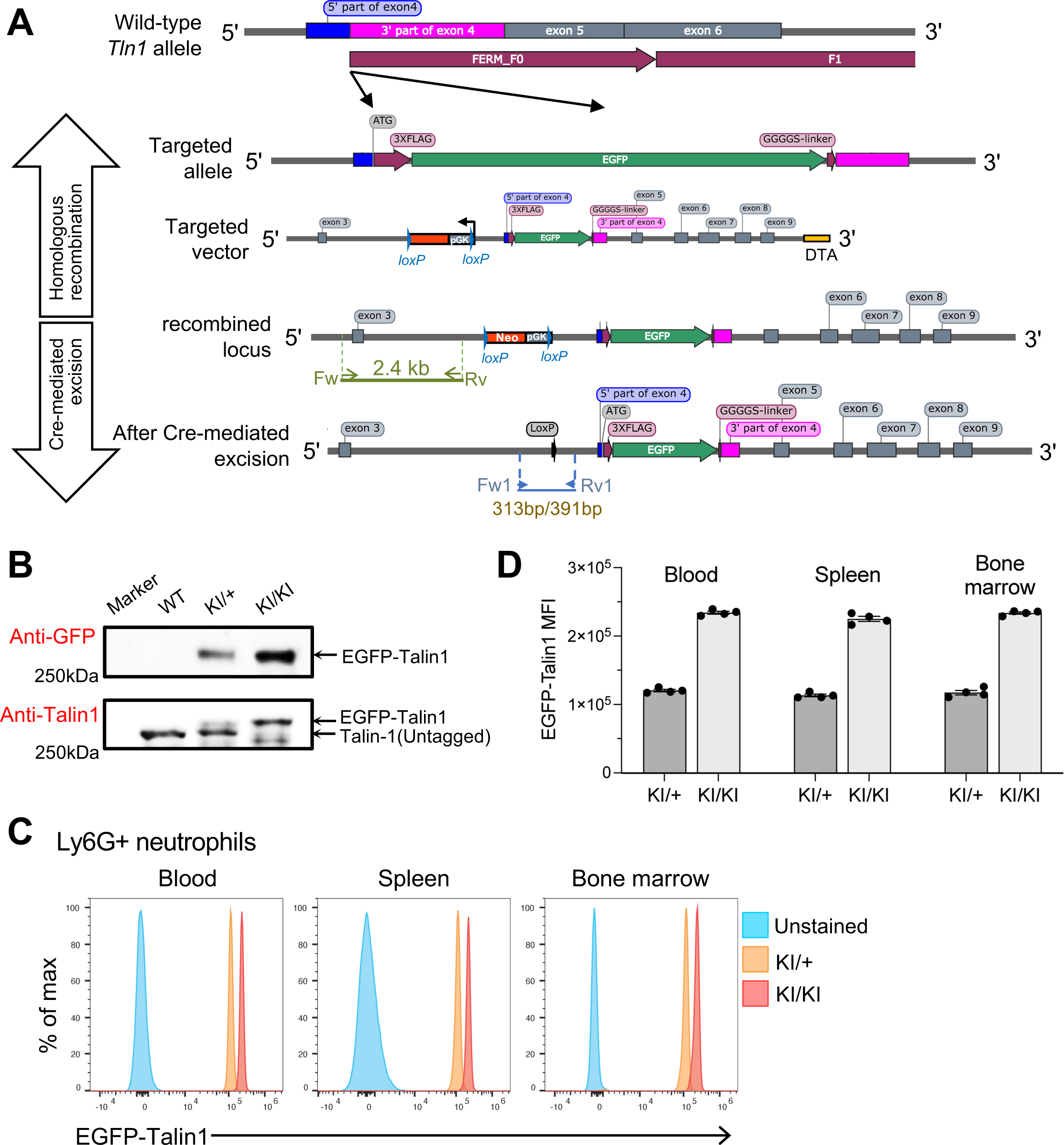
Generation and characterization of EGFP-talin1 knock-in mice. **(A)** Schematic of the targeting strategy used to generate EGFP-talin1 knock-in (KI) mice. The EGFP tag was inserted at the N-terminus of talin-1 with a flexible linker. A floxed selection cassette was removed by Cre-mediated recombination to generate the final knock-in allele. Positions of primer used for homologous recombination in ES cells and for genotyping are indicated. **(B)** Representative immunoblots of neutrophil lysates probed with anti-talin1 and anti-GFP antibodies. EGFP-talin1 migrates at the expected molecular weight (∼ 270 kDa). **(C)** Representative flow cytometry histograms showing EGFP fluorescence in Ly6G+ neutrophils isolated from blood, spleen and bone marrow of WT, KI (KI/+), and KI (KI/KI) mice. **(D)** Quantification of EGFP-talin1 median fluorescence intensity (MFI) in neutrophils from the indicated tissues and genotypes. Data are shown as mean± SEM. Each dot represents an individual mouse. Surface expression of β2 integrins and flow cytometry gating strategies are shown in Fig. S1.

Following arrest, neutrophils crawl along the luminal surface of venules in search of permissive sites for transmigration ^37,38^. The distribution of talin-1 during this intraluminal crawling phase is unknown. Neutrophil transendothelial migration (TEM) occurs through paracellular routes, between endothelial junctions ^39,40^ or transcellular routes, directly through the body of an endothelial cell ^37^. Talin-1 distribution during TEM also has not been investigated. After crossing the endothelium, neutrophils flatten into a “pancake-like” morphology within the subendothelial space ^41^, and subsequently breach the basement membrane at discrete low-expression sites or “portals” ^42^. Talin-1 localization during these late stages of extravasation is also unknown.

Following transmigration through the endothelium and basement membrane, neutrophils begin their interstitial migration while their uropod initially remains tethered to the venular wall, resulting in its marked elongation ^43,44^. The distribution of talin-1 during this phase also remains unknown. Detachment of the uropod is mediated by non-muscle myosin II (type II myosin)-dependent contractility ^44,45^, after which the neutrophil repolarizes ^46^, presumably in response to local chemokine gradients ^47,48^. Once fully polarized, neutrophils exhibit a broad leading edge or lamellipod, a compact cell body, and a narrow, pointed uropod ^44^. Despite this well-characterized morphology, talin-1 localization during post-transmigration polarization and migration remains unknown.

In this study, we generated EGFP-talin1 knock-in mice to visualize the subcellular dynamics of endogenous talin-1 in primary neutrophils in vitro and in vivo. Using total internal reflection fluorescence (TIRF) and intravital microscopy, we tracked talin-1 recruitment during neutrophil rolling, arrest, crawling, transendothelial migration, basement membrane breaching and interstitial migration. Our results reveal distinct patterns of talin-1 localization across these phases and suggest a new role for talin-1 in guiding neutrophil polarization during navigation in tissue environments.

## Materials and Methods

EGFP-talin1 knock-in mice were generated by insertion of a 3xFLAG-EGFP-GGGGS cassette in-frame with the endogenous *Tln1* locus in C57BL/6N-derived embryonic stem cells. Homozygous EGFP-talin1 knock-in mice were generated by intercrossing heterozygous offspring. Fluorophore-conjugated antibodies against Ly6G, CD11a, CD11b, CD18, mAb24, and KIM127 were used for flow cytometry and imaging studies. Recombinant mouse P-selectin-Fc, ICAM-1-Fc, and CXCL1 were used for neutrophil stimulation and adhesion assays. Whole blood, bone marrow, and splenocytes were isolated from mice and stained with fluorophore-conjugated antibodies. Data were acquired on Aurora spectral flow cytometers or a NovoCyte Quanteon flow cytometer and analyzed using FlowJo. For β2 integrin activation assay, bone marrow cells hITGB2 mice^26^ crossed with EGFP-talin1 knock-in mice, as well as control hITGB2 mice. Cells were stimulated with CXCL1, P-selectin-Fc, or Fc crosslinking conditions in the presence of mAb24 and KIM127 antibodies. Neutrophils were identified as live singlet Ly6G+ cells, and β2 integrin activation was quantified by measuring E+H+ cells and fluorescence intensity of mAb24 and KIM127 staining.

Microfluidic perfusion assays were performed as previously described^25,49^. Glass coverslips were coated with recombinant mouse P-selectin-Fc and ICAM-1-Fc. Bone marrow cells from hITGB2 EGFP-talin1 knock-in mice were perfused under 6 dyn/cm^2^ shear stress, and CXCL1 was introduced to induce neutrophil arrest and adhesion.

Neutrophil rolling, arrest, spreading, and migration were visualized by total internal reflection fluorescence microscopy. For intravital microscopy, the mouse cremaster muscle was prepared as previously described ^50,51^. EGFP-talin1 knock-in mice were anaesthetized, and neutrophils were labelled in vivo using Ly6G-Alexa Fluor 647 antibody. Imaging was performed on a NiKon AX R multiphoton microscope using simultaneous dual-channel acquisition. Image analysis was performed using Fiji (ImageJ) and GraphPad Prism 10. Neutrophil rolling, arrest, crawling, transendothelial migration, uropod detachment, and interstitial migration were manually tracked.

Fluorescence intensity profiles were extracted across the cell using rectangular ROIs aligned along the front-to-rear axis, define by the direction of migration. Statistics analyses were performed using GraphPad Prism 10. Comparisons among multiple groups were performed using two-way ANOV with appropriate post hoc multiple-comparison tests. Unpaired two-tailed Mann-Whitney tests were used for comparisons between two independent groups or for each condition, as indicated. Data are presented as mean ± SEM. Statistical significance was defined as *P* < 0.05. Further details are provided in the Supplemental Methods.

## Results

### EGFP-talin1 knock-in mice enable visualization of talin-1 expression without altering basal hematological or β2 integrins expression profiles

To enable real-time visualization of talin-1 dynamics in neutrophils, we generated EGFP-talin1 knock-in mice by inserting EGFP in-frame into exon 4 of the mouse *Tln1* locus (Figure 1A). The targeting construct included a floxed neomycin resistance cassette, which was excised following recombination, resulting in expression of EGFP-talin1 fused to the N-terminal F0 domain via a flexible GGGGS linker (Figure 1 A). Both heterozygous (EGFP-Talin1 KI, KI/+) and homozygous mice (EGFP-Talin1 KI, KI/KI) were viable and fertile, with no obvious spontaneous phenotype (Table S1-3).

Western blot analysis confirmed expression of full-length EGFP-talin1 fusion protein at the expected molecular weight (∼270 kDa) (Figure 1B). Flow cytometry further demonstrated robust EGFP fluorescence in neutrophils from knock-in mice, consistent with gene dosage-dependent expression (Figure 1C and D). EGFP-talin1 expression was comparable across neutrophils isolated from blood, bone marrow and spleen (Figure 1C and D). EGFP-talin1 expression was also detected across multiple immune cell populations (Supplementary Figure 4).

To determine whether insertion of EGFP affected hematopoiesis, complete blood counts were performed. No significant differences were observed in neutrophils, lymphocytes, or other blood cell populations across genotypes (Supplementary Figure 2), indicating normal hematological homeostasis.

We next assessed whether EGFP tagging altered integrin expression. Surface levels of β2 integrin (CD18) and its associated α subunits CD11a and CD11b on neutrophils were comparable between wild-type and knock-in mice (Supplementary Figure 3), indicating that EGFP insertion does not affect integrin surface expression. Together, these data establish EGFP-talin1 knock-in mice as a suitable model for studying talin-1 dynamics in neutrophils.

### EGFP-talin1 preserves β2 integrin activation and adhesion under stimulation

To directly assess β2 integrin activation, EGFP-talin1 knock-in mice were crossed with humanized *hITGB2* mice ^26^, enabling detection of integrin conformation using KIM127 (E+, extended) and mAb24 (H+, high-affinity) antibodies.

Under basal conditions (PBS), neutrophils from both control and EGFP-talin KI mice showed minimal binding of KIM127 and mAb24 (Figure 2A). Stimulation with CXCL1 alone or with P-selectin, or P-selectin-Fc induced β2 integrin activation, as reflected by increased KIM127 and mAb24 signals and the appearance of E+H+ neutrophils (Figure 2A). Crossing PSG1 with P-selectin-Fc and anti-human Fc greatly enhanced the proportion of E+H+ neutrophils by 3-4 fold (Figure 2A right panels). Across all stimulation conditions, the percentage of E+H+ cells, as well as mAb24 and KIM127 fluorescence intensities, were comparable between EGFP-talin1 KI and control neutrophils (Figure 2B), suggesting that EGFP-talin1 supports integrin activation as well as wild-type talin-1. To determine whether preserved integrin activation translates into normal adhesion function, neutrophil behavior under flow was examined. EGFP-talin1 KI neutrophils showed rolling velocities and proportions of firmly adherent cells comparable to controls (Figure 2C), indicating intact β2 integrin-dependent adhesion under shear flow.

**Figure 2.**
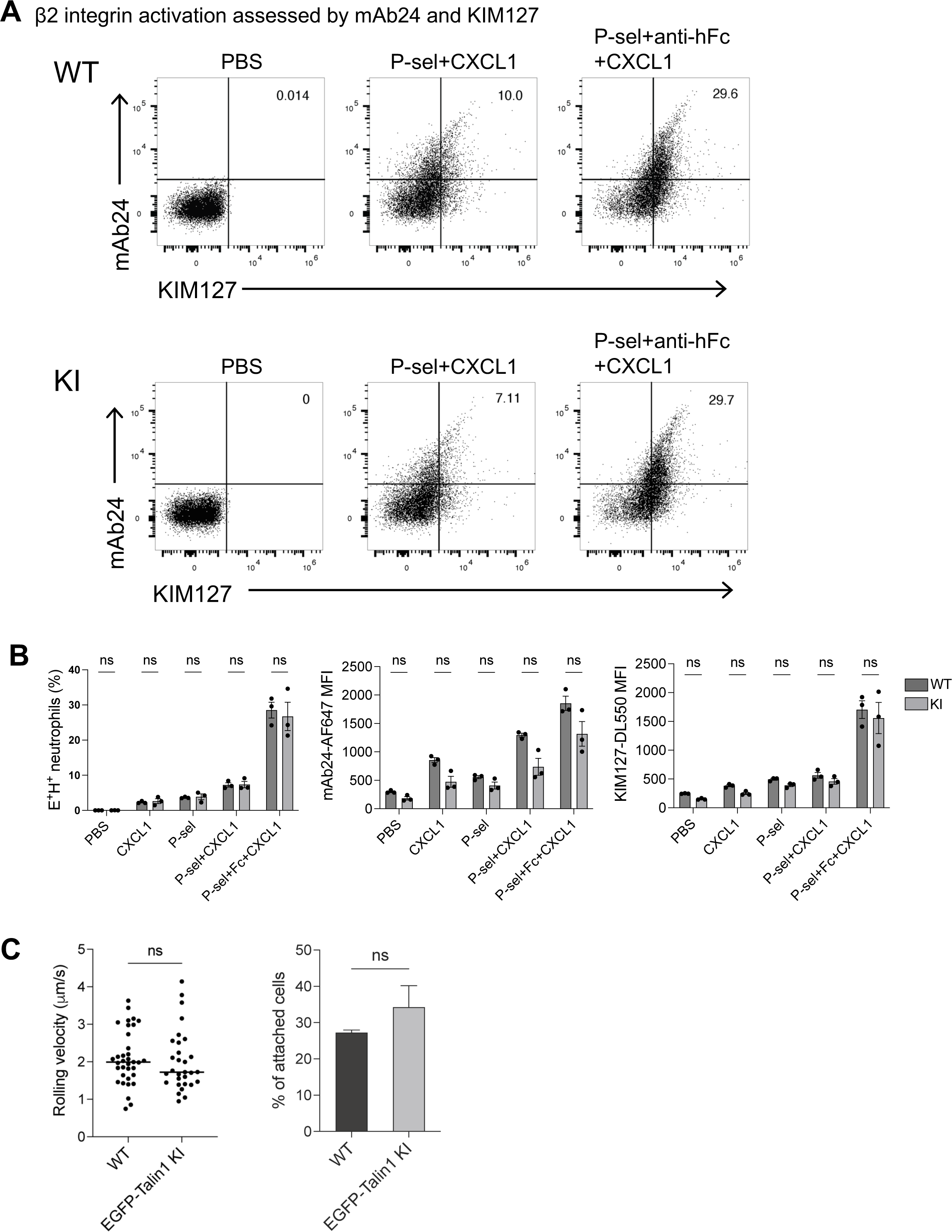
EGFP-talin1 knock-in preserves β2 integrin activation in neutrophils under stimulation. Neutrophils from bone marrow of humanized β2 integrin (hITGB2) mice crossed with EGFP-Talin1 knock-in mice were used to assess β2 integrin activation. PBS, phosphate-buffered saline; CXCL1, C-X-C motif chemokine ligand 1; P-sel, P-selectin; Fc, anti-human Fc antibody used for integrin crosslinking. **(A)** Representative flow cytometry plots showing β2 integrin activation in neutrophils assessed by mAb24 (H+, high-affinity conformation) and KIM127 (E+, extended conformation) under the indicated stimulation conditions. **(B)** Quantification of β2 integrin activation. Left: percentage of E+H+ neutrophils. Middle: mAb24 median fluorescence intensity (MFI). Right: KIM127 MFI. Comparisons between WT and KI neutrophils were performed for each stimulation condition. Each dot represents an individual mouse. **(C)** Neutrophils rolling velocity (left) and percentage of attached cells (right) under flow conditions, reflecting β2 integrin-dependent neutrophil adhesion. Data are shown as mean ± SEM. Statistical significance was determined by unpaired two-tail Mann-Whitney test for each condition (B) or for two-group comparisons (C). ns, not significant.

Together, these data demonstrate that EGFP-talin1 knock-in preserves both β2 integrin activation and downstream adhesive functions in neutrophils.

### Talin-1 accumulates prior to β2 integrin activation during neutrophil arrest under flow

To define the spatiotemporal dynamics of talin-1 during neutrophil rolling and arrest, we performed quantitative dynamic footprinting (qDF) using TIRF imaging under physiological shear flow (6 dyn/cm^2^) ^49^. Neutrophils isolated from EGFP-talin1 knock-in mice were imaged as they transitioned from rolling to arrest to initial spreading.

During rolling, EGFP-talin1 fluorescence within the TIRF field was low, indicating that talin-1 was mainly in the center of the cell and not within ∼100 nm from the membrane. Notably, talin-1 signal began to increase progressively tens of seconds (approximately 20-70 s) prior to arrest and continued to rise at the onset of arrest and early spreading (Figure 3A; and Supplementary Video 1). Quantification of normalized integrated fluorescence intensity confirmed a gradual increase in talin-1 accumulation during rolling, followed by a further increase upon arrest and initial spreading (Figure 3B).

**Figure 3.**
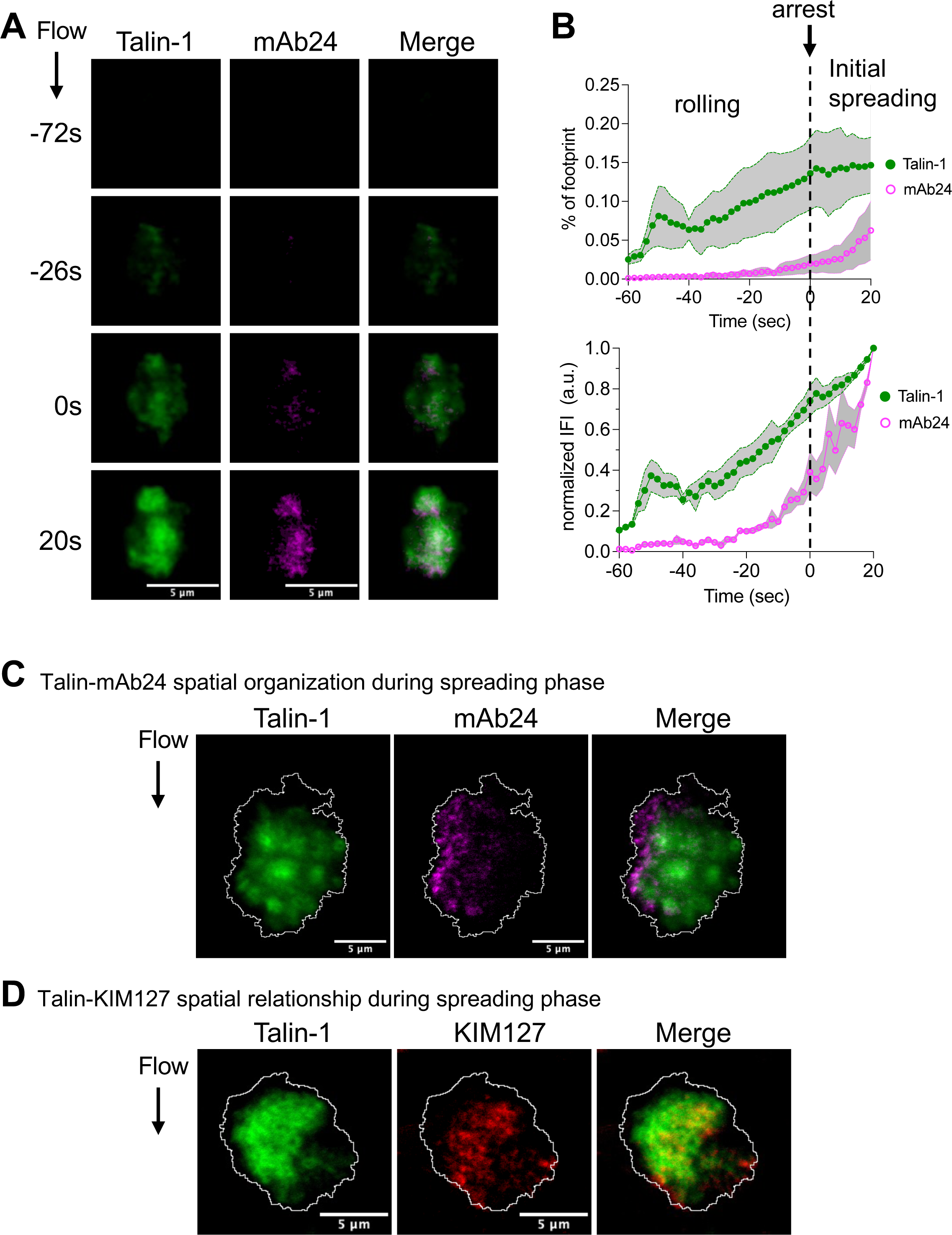
Talin-1 accumulates prior to β 2 integrin activation during neutrophils arrest. Bone marrow neutrophils from humanized β2 integrin (hITGB2) mice crossed with EGFP-Talin1 knock-in mice were labelled with Ly6G-AF647 and perfused over P-selectin and ICAM-1 coated coverslips under shear flow in the presence of CXCL1 to induce arrest. Imaging was performed using quantitative dynamic footprint (qDF) microscopy. **(A)** Representative time-lapse images showing Talin-1 (EGFP) and β2 integrin activation (mAb24) during neutrophil arrest and spreading. Time is indicated relative to the onset of arrest (0 s). Scale bar, 5 μm. See Supplementary video 1. **(B)** Quantification of talin-1 and mAb24 signal over time relative to the onset of arrest. Top: fraction of signal localized with the cell footprint. Bottom: normalized integrated fluorescence intensity (IFI). Solid lines indicate the mean and shaded area represent SEM. n = 3 cells. The dashed line indicates the onset of arrest (0 s). **(C)** Talin1-mAb24 spatial organization. Representative images showing the spatial organization of talin-1 relative to β2 integrin high-affinity conformation (mAb24) during the spreading phase. Cell boundaries are outlined. Scale bar, 5 μm. **(D)** Talin1-KIM127 spatial relationship. Representative images illustrating the spatial relationship between talin-1 and β2 integrin extended conformation (KIM127) during the spreading phase. Cell boundaries are outlined. Scale bar, 5 μm. See Supplementary video 2.

In contrast, high affinity β2 integrin detected by mAb24 displayed a delayed activation profile. mAb24 signal remained minimal during rolling and became detectable only shortly before arrest, followed by a rapid increase after arrest (Figure 3 B), consistent with previous observations in HL60 cells ^25^. Thus, talin-1 recruitment precedes the appearance of high-affinity β2 integrins, indicating that talin-1 membrane association occurs before full integrin activation. Notably, a portion of the mAb24 signal appeared after arrest, consistent with potential outside-in signaling ^52^.

Spatial analysis during spreading further revealed distinct organization of talin-1 and activated integrins. Talin-1 showed a broad distribution across the cell footprint, whereas mAb24 signal appeared toward the rear of the neutrophils in discrete regions (Figure 3C; Supplementary Video 2A). Similarly, KIM127 signal showed partial overlap with talin-1 but was sparser than EGFP signal (Figure 3D; Supplementary Video 2B). A similar spatial pattern was observed in additional cells, including the representative example shown in Supplementary Figure 5. These observations suggest that talin-1 accumulation and integrin conformational activation are temporally and spatially coordinated but not identical processes.

### Talin-1 forms dynamic discrete accumulation sites and redistribution along the front-to-rear axis during neutrophil migration

Following arrest and spreading, a subset of neutrophils initiated polarization and migration under flow. To characterize talin-1 dynamics during this migration, we analyzed EGFP-talin1 distribution in migrating cells using time-lapse TIRF microscopy.

In migrating neutrophils, talin-1 initially showed a relatively broad distribution across the cell footprint but progressively reorganized into discrete accumulation sites that dynamically redistributed over time (Figure 4A; Supplementary Video 3; Supplementary Figure 6). These talin1-enriched regions were not static; instead, they shifted positions as the cell migrated, indicating continuous remodeling of talin-1 localization during cell movement.

**Figure 4.**
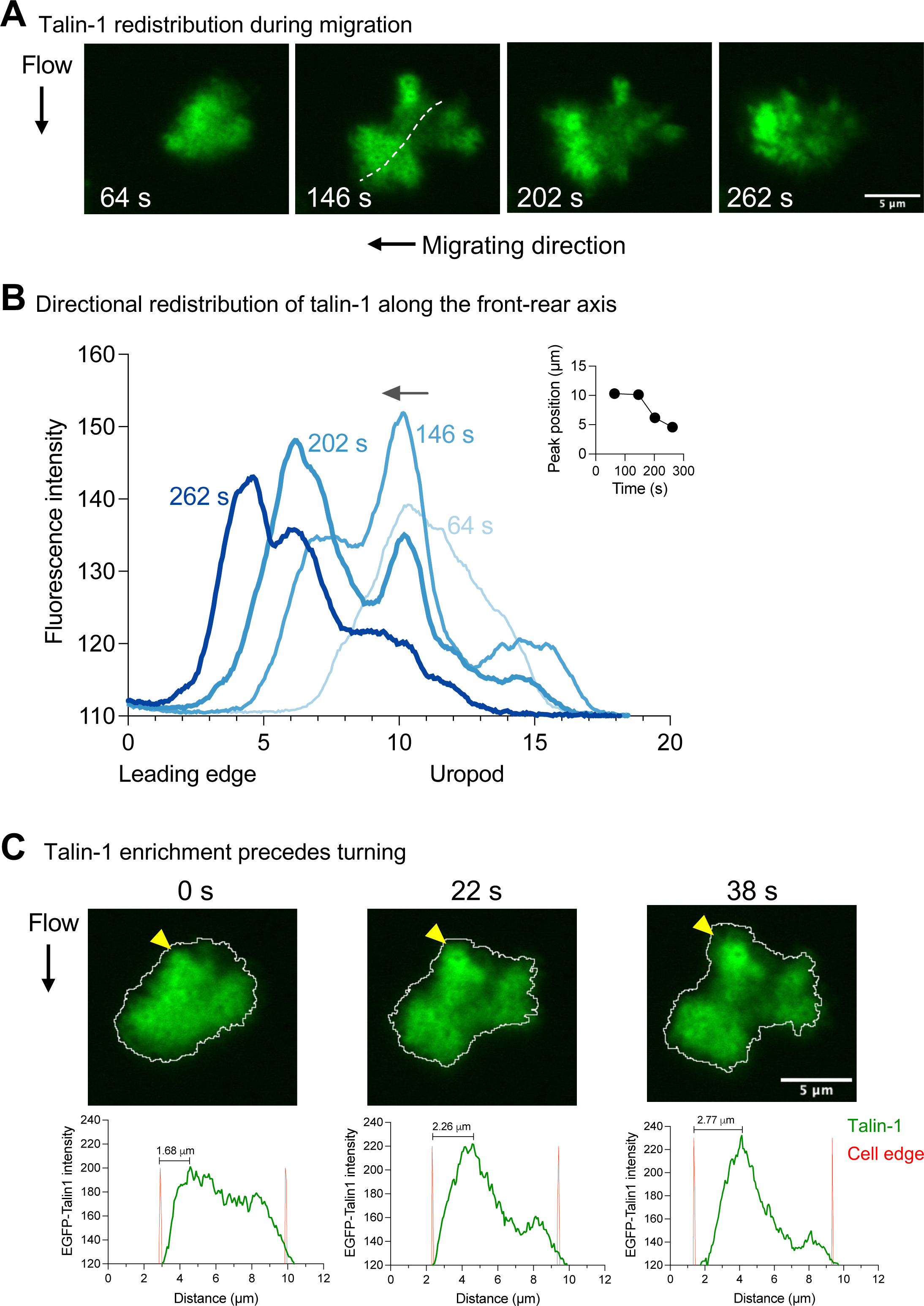
Talin-1 shows dynamic discrete accumulation sites and redistributes along the front-to-rear axis during neutrophil migration. **(A)** Representative time-lapse TIRF images showing directional redistribution of EGFP-talin1 during neutrophil migration under flow. Talin-1 signal is initially broadly distributed and subsequently concentrates into discrete accumulation sites that reorganize over time. Migration direction is indicated. Scale bar, 5 μm. See also Supplementary video 3. **(B)** Quantification of talin-1 distribution along the front-to-rear axis at the indicated time points. Fluorescence intensity profiles were obtained from regions of interest (ROI) spanning the whole cell. Intensity profiles reveal distinct peaks corresponding to talin-1 accumulation sites, which progressively shift toward the leading edge over time. Inset, perk position plotted over time. **(C)** Line-scan analysis of EGFP-talin1 intensity across the cell footprint. Talin-1 enrichment precedes membrane extension toward the turning direction and is followed by a progressive spatial separation from the advancing cell edge. Red line indicate cell boundaries defined from the Ly6G outline. Distance from talin-1 enrichment to the advancing cell edge is indicated.

To quantify these dynamics, we measured fluorescence intensity along the front-to-rear axis using regions of interest spanning the entire cell. Intensity profiles suggested distinct peaks corresponding to talin-1 accumulation sites at each time point (Figure 4B). Notably, the positions of these peaks progressively shifted toward the leading edge over time, consistent with directional redistribution of talin-1 during migration (Figure 4B, inset). Together, these data demonstrate that talin-1 forms dynamic, discrete accumulation sites. Talin-1 localization toward the leading edge appears to determine the direction of migration. To test this hypothesis, we measured EGFP intensity over time relative to motion of the leading edge over time (Figure 4C; Supplementary Figure 7), suggesting a spatially and temporally coordinated redistribution of talin-1 along the front-to-rear axis.

### Stage-specific redistribution of talin-1 during neutrophil extravasation in vivo

To define talin-1 dynamics during neutrophil extravasation in vivo, we performed intravital imaging of the cremaster microcirculation in EGFP-talin1 knock-in mice. During intravascular crawling, talin-1 was distributed along the cell periphery, with enrichment at the front and at the membrane facing the endothelium (Figure 5A). These talin1-enriched regions dynamically reorganized during cell movement, consistent with active adhesion remodeling.

**Figure 5.**
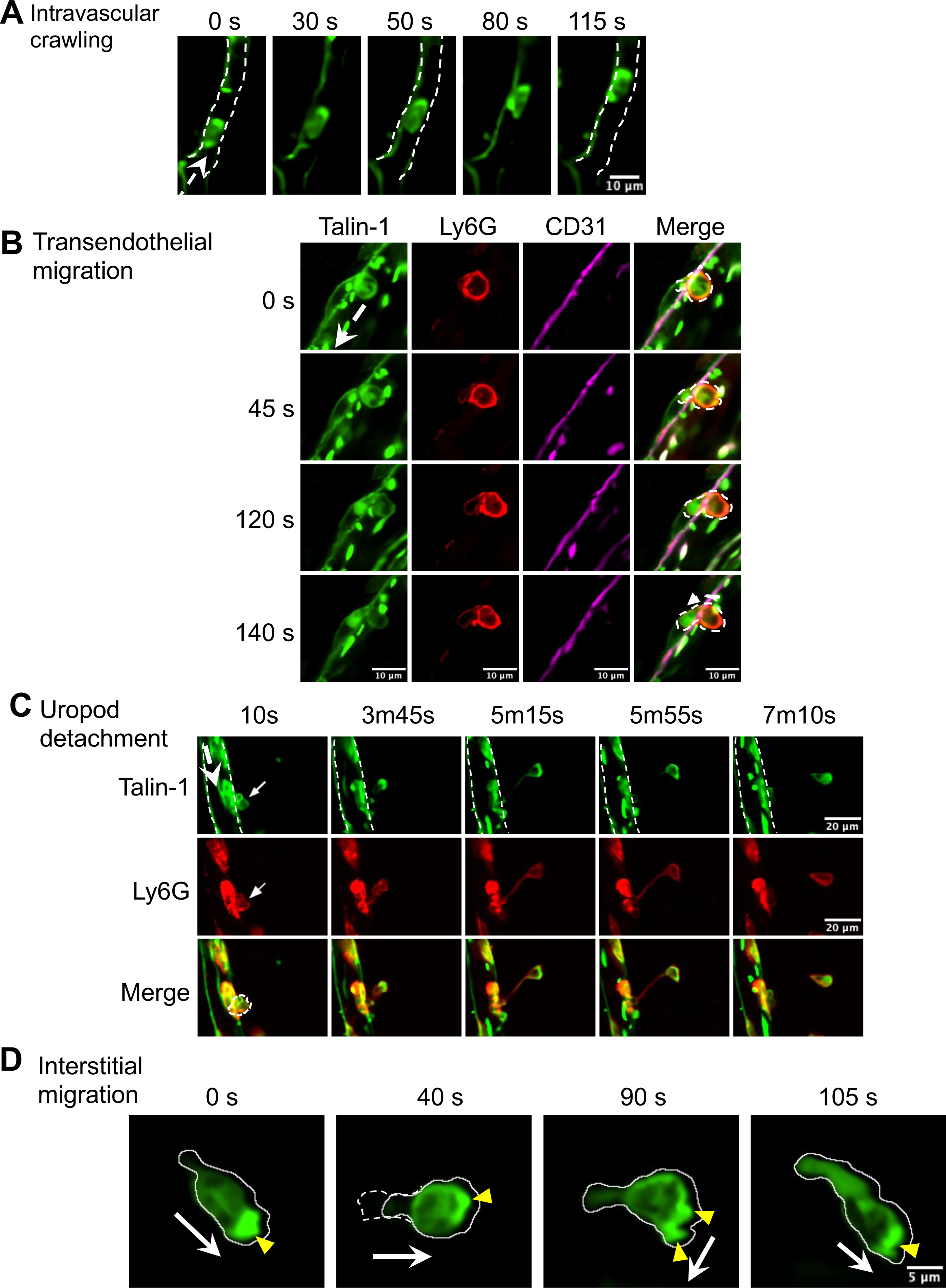
Stage-specific redistribution of talin-1 during neutrophil extravasation. Representative intravital images showing EGFP-talin1 (green) and Ly6G (red) during distinct stages of neutrophil extravasation. **(A)** Intravascular crawling. Neutrophils migrate along the endothelial surface under blood flow, with talin-1 signal enriched at the discrete sites along the cell periphery. Dashed lines outline the vessel wall. Time stamps are indicated. Scale bar, 10 μm. **(B)** Transendogthelial migration. EGFP-talin1 (green), Ly6G (red), and endothelial CD31 (magenta) are shown. Talin-1 accumulates at the neutrophil-endothelial interface during diapedesis, with enrichment at the leading edge and at the sites of endothelial contact. Dashed outlines indicate the cell boundary. Scale bar, 10 μm. **(C)** Uropod detachment. Talin-1 shows front-enriched redistribution as the neutrophil exits the vessel and the rear remains tethered prior to detachment. Ly6G (red) marks the neutrophil. Dashed outlines indicated the cell boundary. Scale bar, 20 μm. **(D)** Representative intravital imaging of a neutrophil undergoing directional turning during interstitial migration. Time stamps indicate seconds. White arrows denote the direction of cell movement, and yellow arrowheads indicate regions of talin-1 enrichment. At 0 s, talin-1 is enriched at the leading edge. By 40 s, the previous position of the uropod is indicated by a dashed outline, illustrating rear retraction. During turning, transient dual talin-1 enrichment sites are observed, followed by the emergence of a dominant talin-1 cluster that aligns with the new direction of migration.

As neutrophils initiated transendothelial migration, talin-1 accumulated at the neutrophil-endothelial interface, with enrichment at the leading edge and at sites of endothelial contact. Talin-1 remained dynamically organized as cells traversed the endothelial barrier (Figure 5 B).

Following diapedesis, talin-1 remained enriched toward the leading edge as neutrophils exited the vessel, while the uropod of the cell remained transiently tethered prior to complete uropod detachment, consistent with previous reports ^53^ (Figure 5C; Supplementary Video 4). After extravasation, neutrophils continued to migrate within the interstitial tissue (Supplementary Video 5). Just like in the flow chamber, the accumulation of talin-1 near the leading edge preceded each turn (Figure 5D).

## Discussion

Talin-1 is a key regulator of integrin activation, linking integrins to the actin cytoskeleton and inducing conformation changes that promote high-affinity binding. While its role in integrin-mediated adhesion has been well characterized in vitro and in non-leukocyte systems ^35,54,55^, its spatiotemporal dynamics during neutrophil recruitment and migration in vivo have remained poorly understood. In this study, we developed EGFP-talin1 knock-in mice to enable direct visualization of talin-1 behavior under physiological conditions. Using TIRF and intravital microscopy, we show that talin-1 is dynamically recruited during each stage of the neutrophil adhesion cascade, preceding arrest, redistribution during luminal crawling and transmigration, and remaining polarized during interstitial migration. These findings provide new insight into how talin-1 coordinates integrin function with cytoskeletal remodeling to support leukocyte trafficking.

Our in vitro flow chamber experiments reveal that talin-1 accumulates in the neutrophil footprint during slow rolling and increases further at the onset of arrest. This temporal pattern suggests that talin-1 recruitment precedes full integrin activation, consistent with a model in which talin-1 engagement represents an early step in integrin activation ^35,56^. Importantly, the delayed emergence of the mAb24 epitope highlights a temporal separation between talin-1 membrane association and high-affinity β2 integrin formation. These findings indicate that talin-1 accumulation and integrin conformational activation are coordinated but not identical processes, consistent with talin-1’s proposed role in promoting integrin activation ^57–59^, but suggests that talin-1 accumulation alone is not sufficient to induce the mAb24 epitope. One candidate molecule that is known to be required for inducing the mAb24 epitope is kindlin-3 ^60^.

In migrating neutrophils, talin-1 did not remain uniformly distributed but instead reorganized into discrete accumulation sites that dynamically redistributed over time. quantitative analysis showed that these sites formed distinct intensity peaks along the cell axis, which progressively shifted toward the leading edge during migration. This behavior suggests that talin-1 localization is continuously remodeled in a directional manner.

Intravital imaging further demonstrated that talin-1 remains highly dynamic after arrest. During luminal crawling, talin-1 was distributed along the cell periphery with enrichment at discrete sites, consistent with continuous adhesion remodeling as neutrophils migrate along the endothelium. These findings align with previous studies identifying LFA-1, Mac-1 and ICAM-1 interactions as the key mediators of crawling ^53,61,62^ and extend them by showing the spatial dynamics of talin-1 during this process in vivo.

During transendothelial migration, talin-1 accumulated at the neutrophil-endothelial interface and beneath the cell body, consistent with a role in reinforcing integrin-cytoskeleton linkages and generating traction forces required for breaching endothelial junctions ^63,64^. As neutrophils progressed through the endothelium, talin-1 was enriched toward the leading edge. The rear of the cell showed little EGFP signal and remained tethered prior to complete uropod detachment. This process may be influenced by endothelial signals such as CD31, which has been shown to promote uropod disengagement during transmigration ^65^. This spatial asymmetry suggests localized regulation of integrin-cytoskeleton coupling along the cell axis. These observations support a model in which reduced talin-1 association at the rear facilitates uropod release. In the interstitial tissue, talin-1 redistribution remained dynamic during neutrophil migration, as observed in addition recordings. Although interstitial 3D migration is less dependent on integrin-mediated adhesion ^66^ than migration on a 2D surface, the persistence of talin-1 polarity suggests that talin-1 may contribute to front-directed organization even in this context.

Together, these findings support a model in which talin-1 redistribution is tightly coupled to forward progression during neutrophil migration. Rather than acting solely as a static integrin activator, talin-1 undergoes continuous spatial reorganization, forming dynamic accumulation sites that shift toward the leading edge and coordinate adhesion, cytoskeletal engagement, and polarity across distinct migratory stages (Figure 6). In this framework, reduced talin-1 association at the rear may facilitate uropod detachment, whereas front-enriched accumulation supports directional migration.

**Figure 6.**
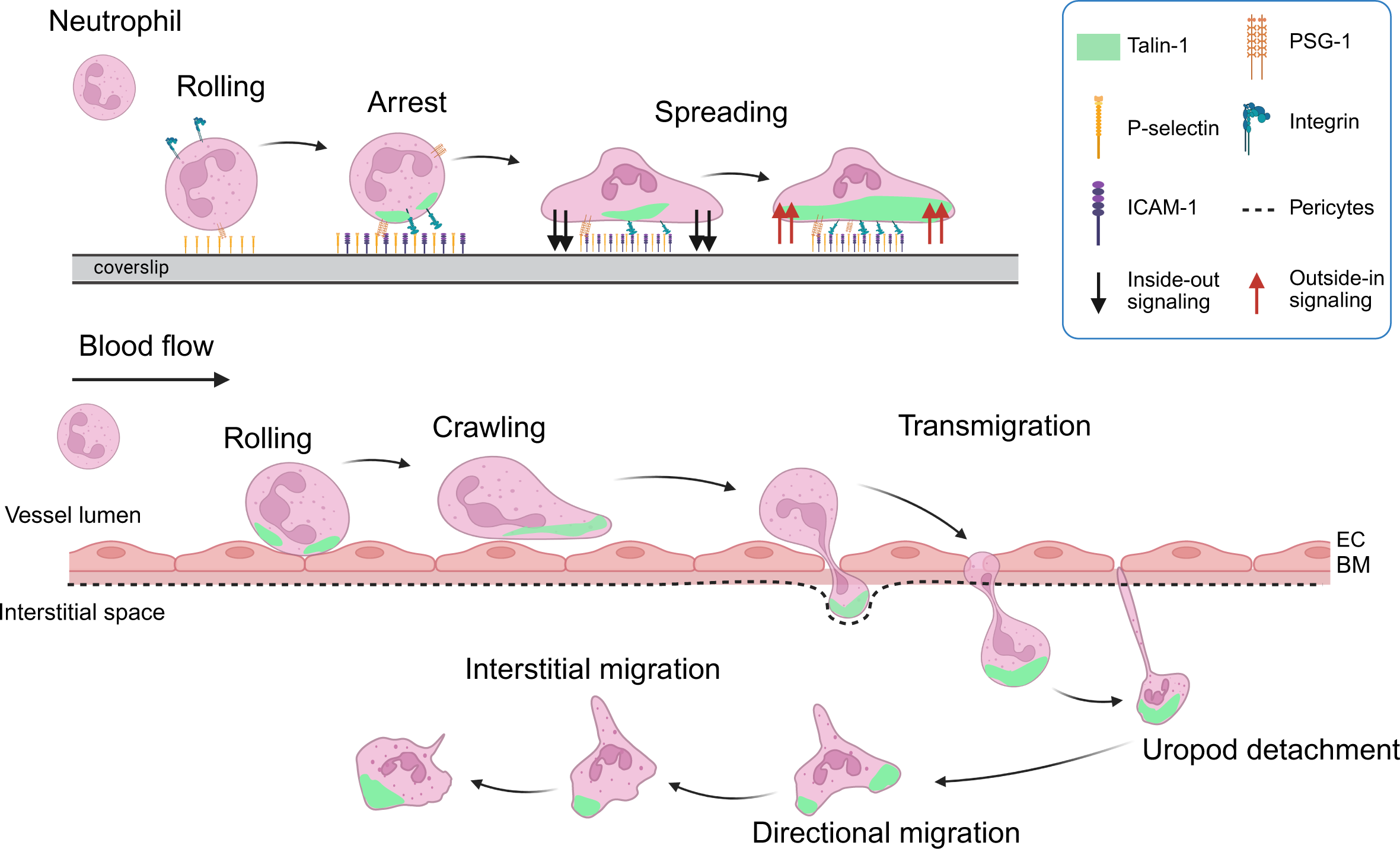
Proposed model of stage-specific talin-1 redistribution during neutrophil recruitment and migration. Schematic illustration summarizing talin-1 localization across the neutrophil adhesion cascade. Talin-1 is recruited to the plasma membrane during rolling and arrest, accumulates at the leading edge during luminal crawling, and redistributes to sites of endothelial contact during transmigration. Following extravasation, talin-1 shows front-enriched distribution during interstitial migration. These stage-specific patterns are consistent with roles for talin-1 in integrin activation and in coordinating adhesion-dependent processes during neutrophil trafficking.

While EGFP-talin1 faithfully supports normal neutrophil function, we cannot exclude the possibility that fluorescent tagging subtly affects talin-1 localization or interactions. Our study also did not address the role of talin-2, which may compensate for or act in parallel with talin-1 in certain immune contexts, although talin-1 is considered the predominant talin isoform in hematopoietic cells^67^.

In conclusion, our study defines the spatiotemporal dynamics of talin-1 during sequential stages of neutrophil trafficking in vivo. We show that talin-1 accumulates prior to arrest, precedes the appearance of the high affinity β2 integrin conformation and is more broadly distributed that high affinity (mAb24+) and extended (KIM127+) β2 integrin molecules. Talin-1 reorganized into dynamic accumulation sites during migration with accumulation toward the front and the neutrophil-endothelial contact site. Talin-1 accumulates near the leading edge, and its peak concentration precedes each turn. Thus, talin-1 spatially determines the directionality of neutrophil migration.

## Acknowledgments

We thank the intravital imaging core and the Georgia Cancer Center Flow and Mass Cytometry Core (RRID: SCR_025747) for their expert support with our research. This work was supported by the Ginsberg Program Project Grant (PPG; P01HL151433) from the National Heart, Lung, and Blood Institute, NIH.

## Authorship Contributions

Y.W. and K.L. designed the experiments. Y.W. performed most of the experiments and data analysis. Q.L. performed TIRF imaging. S.P. conducted western blot experiment and analysis. K.L. supervised the project. Y.W. and K.L. wrote the manuscript. All authors reviewed the data, provided feedback, and revised the manuscript.

