## Supplementary Materials for "Talin-1 determines the direction of primary mouse neutrophils migrating in vivo"

**Expanded Methods**

**Supplementary Tables 1-3**

**Supplementary Figure 1-7**

**Supplementary Video legends**

**Supplementary References**

**Expanded** **Methods**

**Mice**

Wild-type C57BL/6 mice (10-12 weeks, male or female) were purchased from The Jackson Laboratory and maintained under specific pathogen-free conditions at Augusta University (AU). All animal procedures were approved by the AU Institutional Animal Care and Use Committee.

EGFP-talin1 knock-in mice were generated by GenOway by inserting a 3xFLAG-EGFP-GGGGS cassette in-frame with the endogenous start codon (ATG) located in exon 4 of the mouse *Tln1* gene. Homologous recombination was performed in C57BL/6N-derived embryonic stem (ES) cells. The *lox*P-flanked neomycin-resistance cassette was removed by Cre-mediated recombination. Homozygous EGFP-talin1 mice were subsequently generated by intercrossing heterozygous offspring. Expression of the EGFP-talin1 fusion protein was confirmed by western blotting.

**Reagents**

PE-conjugated anti-mouse CD11a antibody (I21/7), BV650-conjugated anti-mouse CD11b antibody (M1/70), and Alexa Fluor 700-, Alexa Fluor 647-, or BV421-conjugated anti-mouse Ly-6G antibody (1A8) were purchased from BioLegend. BUV805-conjugated anti-mouse CD18 antibody (GAME-46) was obtained from BD Biosciences. Live/Dead Fixable Blue Dead Cell stain was purchased from Thermo Fisher Scientific. Anti-human mAb24, a conformation-specific integrin reporter antibody, was purchased from BioLegend (Alexa fluor 647-conjugated) or conjugated in-house to DyLight 550 using the DyLight antibody labelling kit (Thermo Fisher Scientific). Anti-human KIM127 antibody was purified from hybridoma supernatant (University of Virginia) and conjugated in-house to DyLight 550 (Thermo Fisher scientific). Additional reagents included anti-talin1 (8D4, Abcam), anti-GFP (Sigma-Aldrich). protein G, casein blocking buffer, recombinant mouse P-selectin Fc, CXCL1, and ICAM-1 (R&D systems), AffiniPure® F(ab')₂ Fragment Goat Anti-Human IgG Fcγ fragment specific (Jackson ImmunoResearch Laboratories). Neutrophils were enriched using the EasySep Mouse Neutrophil Enrichment Kit (STEMCELL Technologies).

**Flow cytometry**

Mouse whole blood was collected by cardiac puncture. Bone marrow cells were isolated from tibias and splenocytes from spleens. Red blood cells were lysed, and live cells were stained with Live/Dead Fixable Blue viability dye for 30 min at 4°C. Surface markers were labeled by incubation with directly conjugated fluorescent antibodies for 30 min at 4°C. Data were acquired on Aurora spectral flow cytometers (Cytek Biosciences) and analyzed with FlowJo v10.10. Gating strategy is shown in Fig. S1.

**Complete blood counts**

Mouse blood was collected via retro-orbital bleeding into EDTA-coated tubes. Blood counts, including total white blood cells, neutrophils, lymphocytes, red blood cells, platelets, and hemoglobin concentration, were measured using a Hemavet analyzer (Drew Scientific) according to the manufacturer’s instructions. Wild-type, EGFP-tallin1 KI (KI/+), and EGFP-talin1 KI (KI/KI) mice were used.

**β2 integrin activation assay**

Bone marrow cells were isolated from humanized β2 integrin (hITGB2) mice crossed with EGFP-talin1 knock-in mice, as well as control hITGB2 mice. Cells were first stained with BV421-conjugated anti-mouse Ly6G antibody in PSE buffer (PBS supplemented with 2% FBS and 1 mM EDTA) for 30 min at 4°C. EDTA-containing buffer was used during staining to minimize unintended integrin activation. After washing, cells were re-suspended in phenol red-free RPMI 1640 supplemented with 2% human serum albumin for stimulation.

A total of 1×10^6^ cells were stimulated with PBS, CXCL1 (100 ng/ml), P-selectin Fc (5 μg/ml), P-selectin-Fc in the presence of CXCL1, or P-selectin-Fc plus anti-human Fc crosslinking antibody in the presence of CXCL1. During stimulation, cells were incubated with Alexa Fluor 647-conjugated mAb24 (1 μg/ml) and DyLight 550-conjugated KIM127 (1 μg/ml) antibodies for 10 min at room temperature with gentle shaking.

Following stimulation, cells were fixed with 1% paraformaldehyde. Data were acquired on a NoveCyte Quanteon flow cytometer (Agilent) and analyzed with FlowJo v10.10. Neutrophils were gated as live, singlet Ly6G+ cells, and the percentage of E+H+ (KIM127+ mAb24+) cells as well as median fluorescence intensity (MFI) of mAb24 and KIM127 were quantified.

**Microfluidic perfusion assay**

Microfluidic devices were assembled as previously described, with minor modifications^1,2^. Briefly, glass coverslips were coated with recombinant mouse P-selectin-Fc (5 μg/ml) and ICAM-1-Fc (10 μg/ml) for 2 h at room temperature and subsequently blocked with 1% casein for 1 h. Coated coverslips were mounted onto polydimethylsiloxane (PDMS) microfluidic devices using magnetic clamps to form flow channels approximately 29 μm in height and 300 μm in width.

Bone marrow cells were isolated from humanized β2 integrin (hITGB2) EGFP-talin1 knock-in mice and stained with Alexa Fluor 647-conjugated anti-mouse Ly6G antibody. Cells were perfused through the microfluidic channels at a wall shear stress of 6 dyn/cm^2^. CXCL1 (100 ng/ml) was introduced through the inlet to induce neutrophil arrest and adhesion.

Neutrophil behavior, including rolling, arrest, adhesion, spreading, and migration, was recorded by TIRF microscopy and analyzed as described below. All experiments were performed under identical flow and coating conditions.

**Intravital microscopy**

Intravital microscopy of the mouse cremaster muscle microcirculation was performed as previously described ^3,4^. Male EGFP-talin1 knock-in mice were anaesthetized by intraperitoneal injection of xylazine (10 mg/kg) and ketamine hydrochloride (100 mg/kg) and maintained on a heating pad at 37°C. The right jugular vein was cannulated for intravenous administration of antibodies and stimuli. The cremaster was exteriorized, cauterized, and positioned under a coverslip for imaging.

Imaging was performed on a Nikon AX R multiphoton microscope with a 20x/1.00 NA water-immersion objective. Neutrophils were visualized with intravenous Ly6G-Alexa Fluor 647 (5 μg). EGFP-talin1 and Ly6G-AF647 signals were excited using 488 nm and 640 nm lasers, respectively, with emission collected at 500-550 nm (EGFP) and 663-738 nm (Alexa Fluor 647). Images were acquired usingin simultaneous dual-channel resonant scanning.

**Image analysis**

Image data were analyzed using Fiji (ImageJ) and GraphPad Prism 10. Neutrophil behaviors including rolling, arrest, crawling, and transendothelial migration, were manually tracked. Arrest was defined as the first frame in which the cell ceased translational motion and remained stationary for at least 5 seconds.

For fluorescence intensity analysis, regions of interest (ROIs) encompassing the entire cell footprint were defined based on Ly6G signal to delineate cell boundaries. Fluorescence signals were background-subtracted and normalized by min-max scaling.

To quantify talin-1 spatial distribution fluorescence intensity profiles were extracted across the cell using rectangular ROIs aligned along the front-to-rear axis, defined by the direction of migration. Intensity profiles were obtained using the Plot Profile function in Fiji.

Peak positions were determined as the coordinates of maximum fluorescence intensity along the cell axis. Distances between talin-1 intensity peaks and the nearest cell edge were calculated along the line scan axis to quantify spatial separation between talin-1 enrichment and the plasma membrane.

For time-resolved analysis of directional changes, cells undergoing turning events were identified based on changes in migration direction over time. Representative time points were selected to illustrate talin-1 redistribution relative to cell morphology and movement direction.

**Statistics analysis**

Statistical analyses were performed using GraphPad Prism 10. Comparisons among multiple groups were performed using two-way ANOV with appropriate post hoc multiple-comparison tests. Unpaired two-tailed Mann-Whitney tests were used for comparisons between two independent groups or for each condition, as indicated. Data are presented as mean ± SEM. Statistical significance was defined as *P* < 0.05.

**Supplementary Tables**

**Supplementary Table 1. Genotype distribution**

| Breeding  Pair | *Tln1*^+/+^ (n) | *Tln1*^EGFP-Tln1/+^ (n) | *Tln1*^EGFP-Tln1/EGFP-Tln1^ (n) | Total pups | Mendelian χ² Test (p-value) |
| --- | --- | --- | --- | --- | --- |
| Het x Het (Tln1^EGFP-Tln1/+^ × Tln1^EGFP-Tln1/+)^ | 4 | 13 | 11 | 28 | *p* = 0.1618 |

**Supplementary Table 2. Litter size by breeding type**

| Breeding Pair | Litter Size (mean ± SD) | Reference (JAX C57BL/6J) |
| --- | --- | --- |
| Het x Het (Tln1^EGFP-Tln1/+^ × Tln1^EGFP-Tln1/+)^ | 7.3 ± 2.6 | ~6.7 pups/litter |
| Het × Homo (Tln1^EGFP-Tln1/+^ × *Tln1*^EGFP-Tln1/EGFP-Tln1)^ | 6.4 ± 1.8 |  |
| Homo × Homo (*Tln1*^EGFP-Tln1/EGFP-Tln1^ × *Tln1*^EGFP-Tln1/EGFP-Tln1)^ | 7.3 ± 2.5 |  |

**Supplementary Table 3. Body weight by age and sex for homozygous (*Tln1*^EGFP-Tln1/EGFP-Tln1^)**

| Sex | Age | n | Body weight (g, mean ± SD) | JAX reference (g) |
| --- | --- | --- | --- | --- |
| Male | 6 weeks | 10 | 21.5 ± 0.6 | 21.9 ± 1.8 |
|  | 8 weeks | 15 | 22.7 ± 2.1 | 25.0 ± 1.4 |
|  | 12 weeks | 14 | 26.5 ± 1.7 | 28.9 ± 2.0 |
| Female | 6 weeks | 11 | 17.3 ± 1.2 | 18.5 ± 0.9 |
|  | 8 weeks | 11 | 18.8 ± 1.5 | 19.6 ± 1.2 |
|  | 12 weeks | 11 | 21.0 ± 2.8 | 21.9 ± 1.6 |

**Supplementary Figures and Legends**

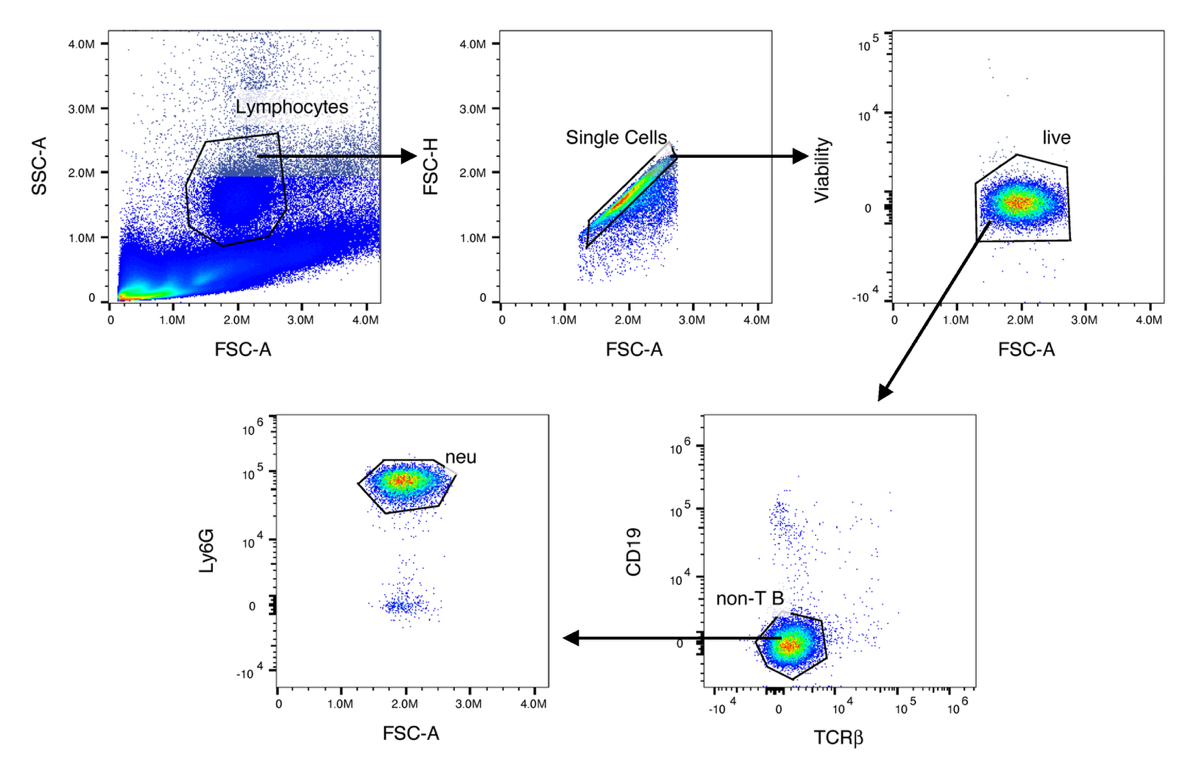

**Supplementary Figure 1.** **Flow cytometry gating strategy for neutrophil and immune cell identification.** Representative flow cytometry plots showing the gating strategy used to identify neutrophils and immune cell populations. Cells were sequentially gated on lymphocytes by forward and side scatter, singlets, and live cells, followed by identification of Ly6G+ neutrophils. Lymphocyte populations were further defined based on CD19 and TCRβ expression.

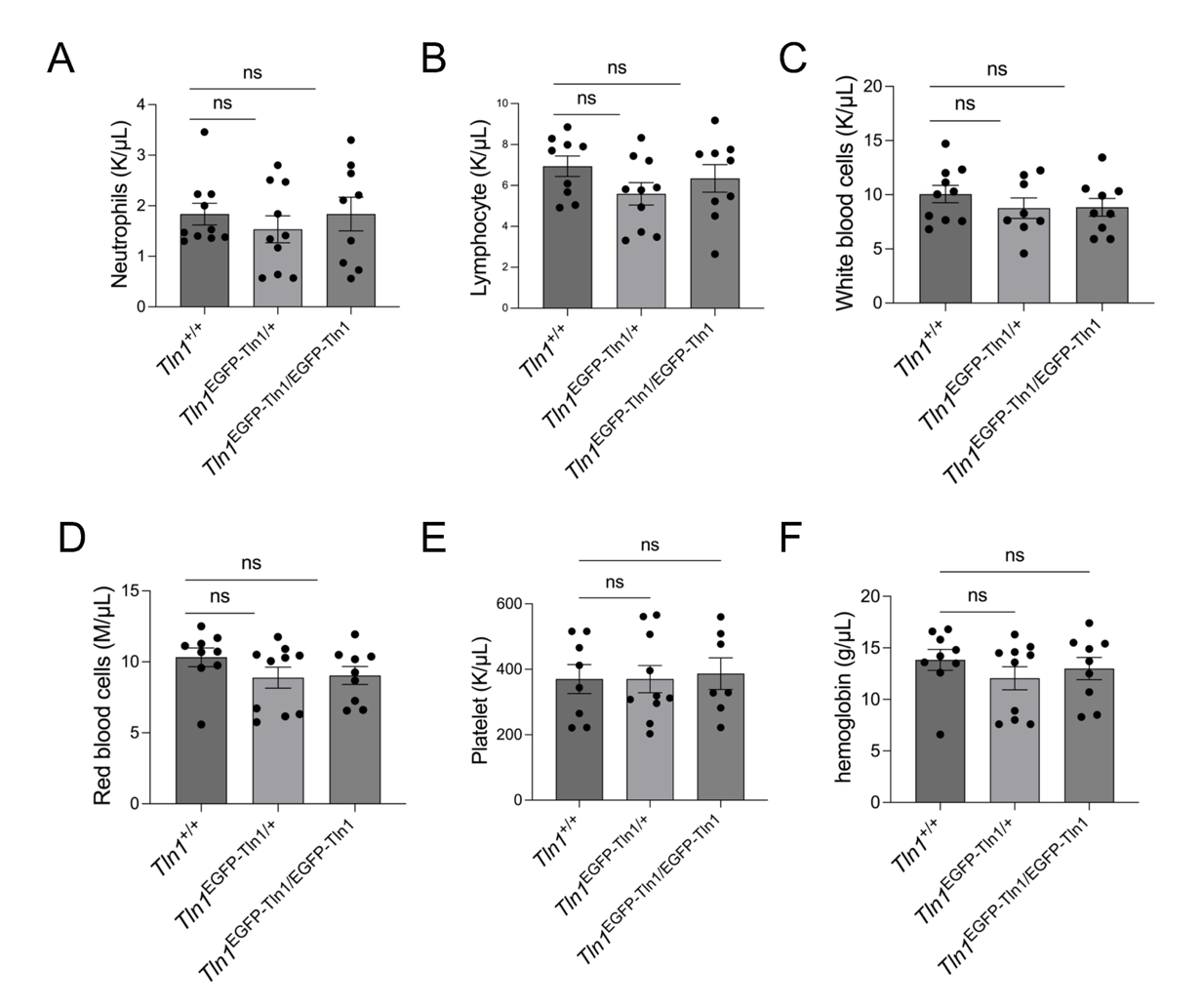

**Supplementary Figure 2**. **Hematological characterization of EGFP-talin1 knock-in mice.** Complete blood counts were performed on WT, EGFP-talin1 KI (KI/+), and EGFP-talin1 KI (KI/KI) mice. Neutrophils (A), lymphocytes (B), total white blood cells (C), red blood cells (D), platelets (E), and hemoglobin levels (F) were quantified using a Hemavet analyzer. Five mice per genotype were measured with technical replicates. Data are presented as mean ± SEM.

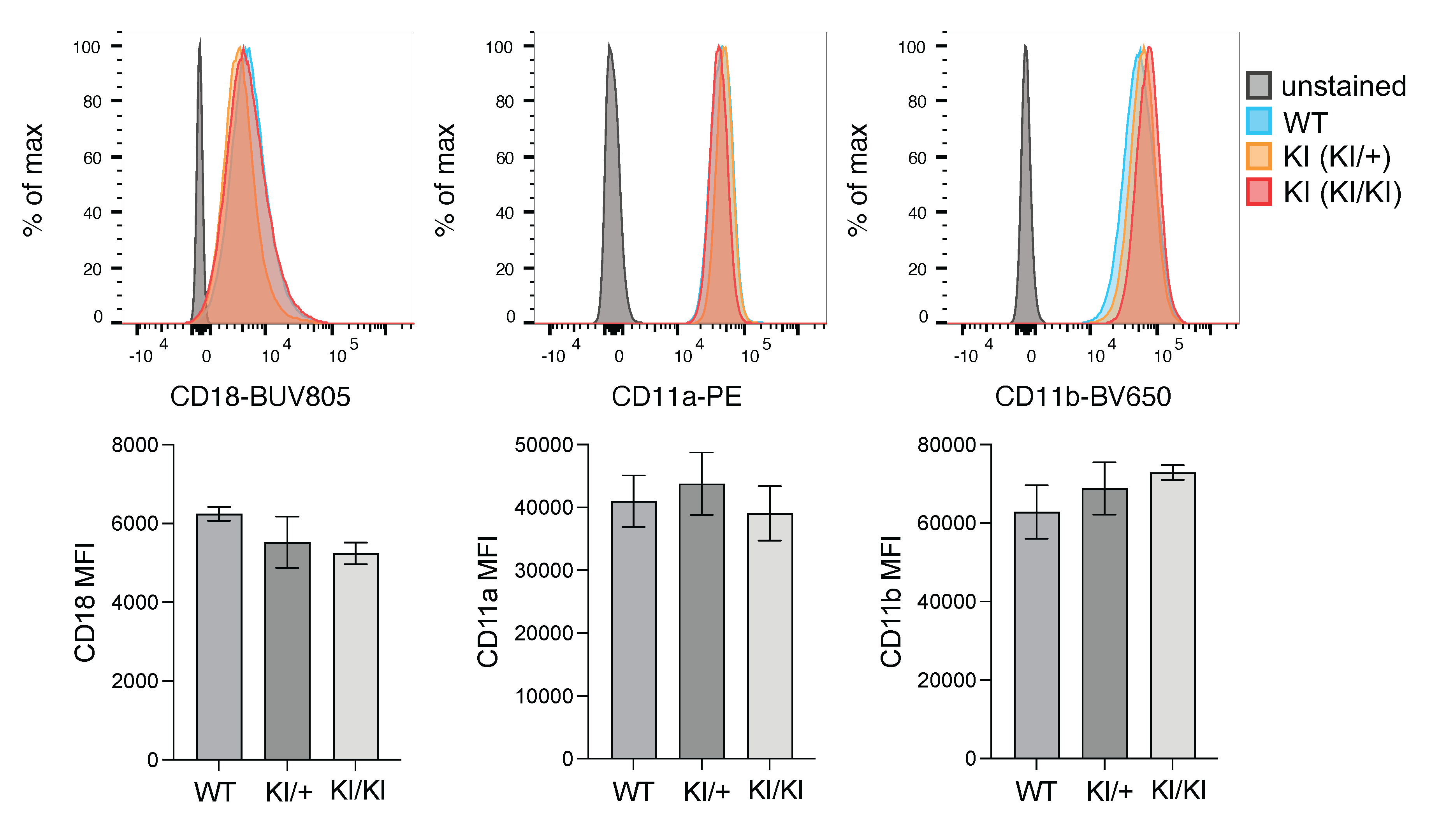

**Supplementary Figure 3**. **Surface expression of β2 integrins in neutrophils from EGFP-talin1 knock-in mice.** Representative flow cytometry histogram (top) and quantification of median fluorescence intensity (MFI; bottom) showing surface expression of CD18, CD11a and CD11b in bone marrow neutrophils from WT, EGFP-Talin1 KI (KI/+), and EGFP-Talin1 KI (KI/KI) mice. Histograms are representative of independent experiments. Bar graphs summarize MFI across mice. n=4 mice. Data are shown as mean ± SEM.

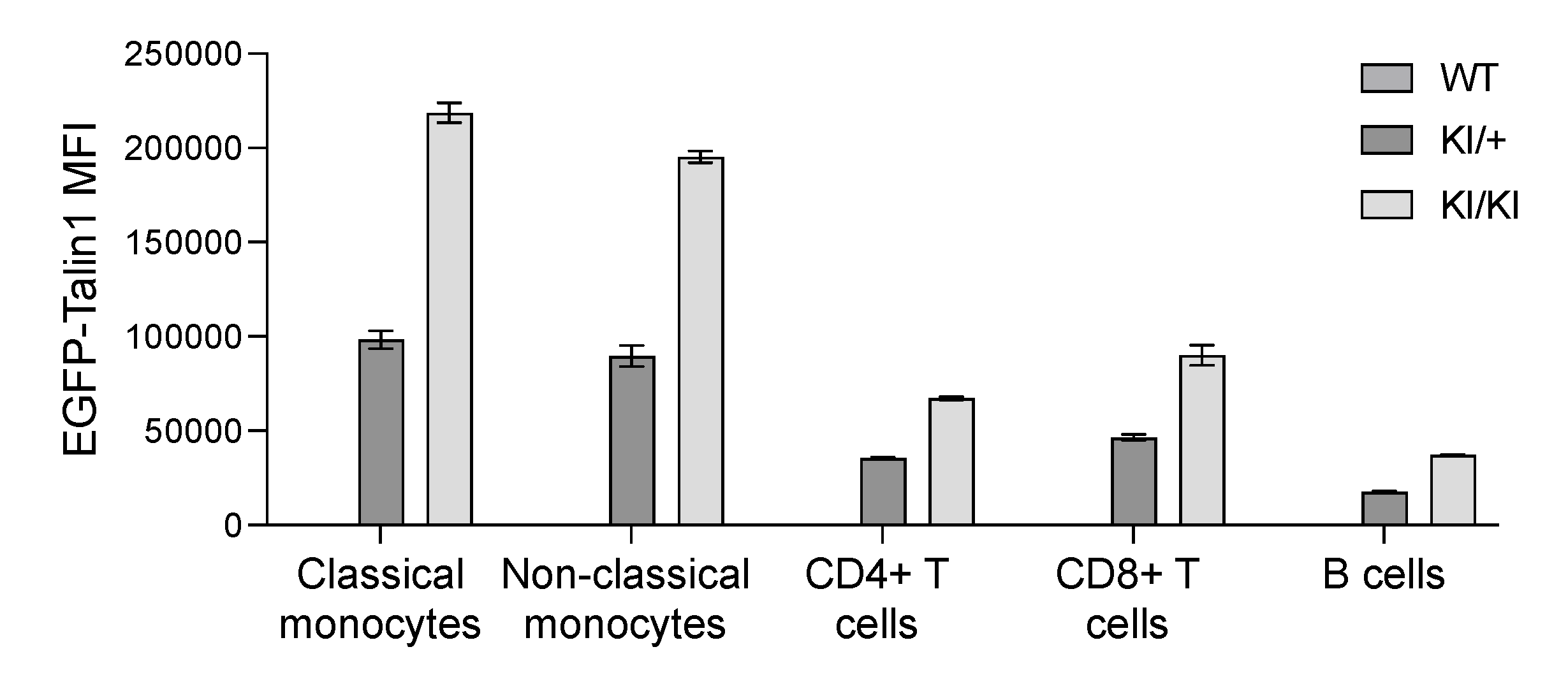

**Supplementary Figure 4. EGFP-talin1 expression across immune cell populations.** Quantification of EGFP-talin1 expression in immune cell populations, including classical monocytes, non-classical monocytes, CD4+ T cells, CD8+ T cells, and B cells. Expression levels are presented as median fluorescence intensity (MFI). n=3 mice. Data are shown as mean ± SEM.

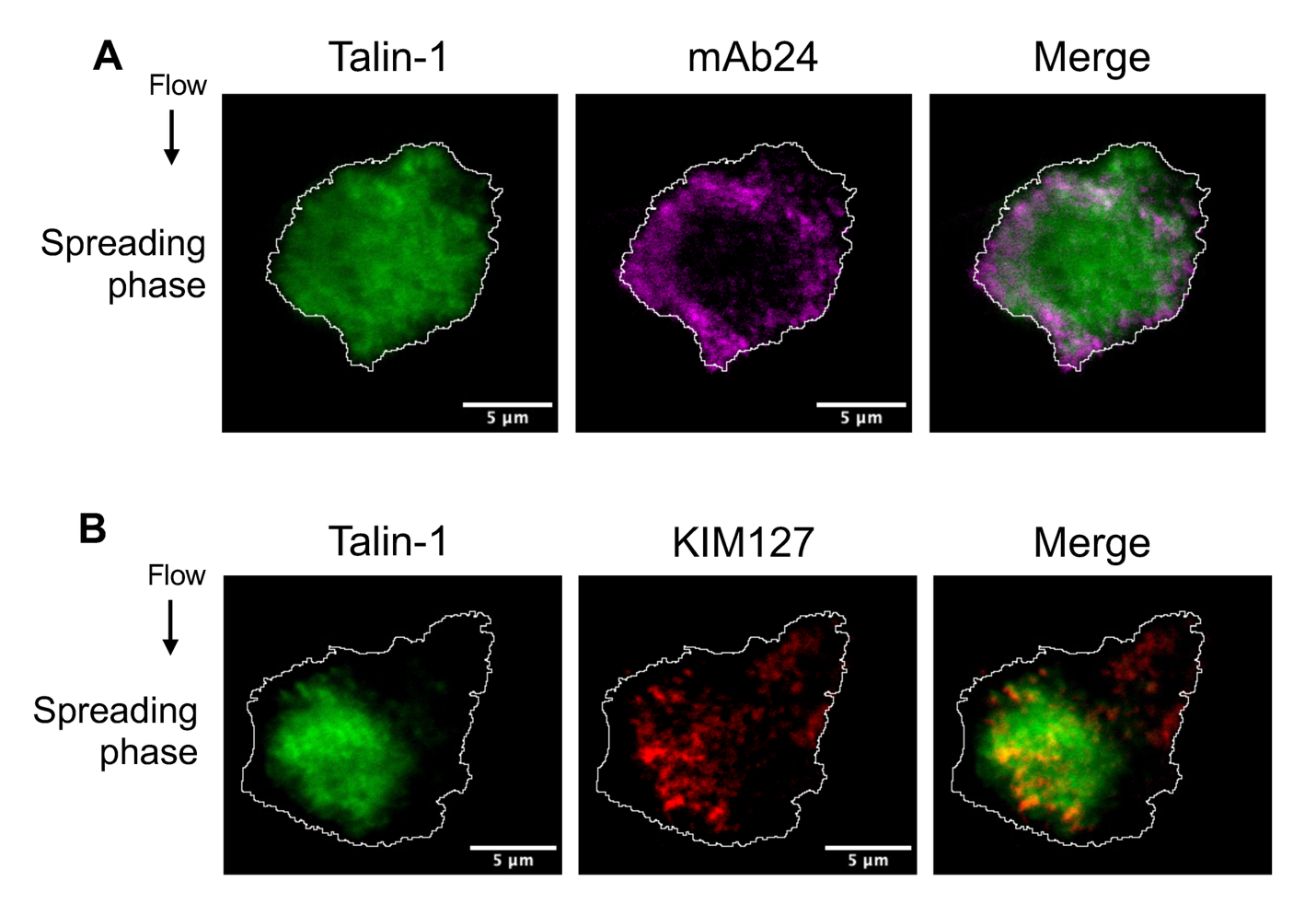

**Supplementary Figure 5.** **Additional representative neutrophils showing talin-1 spatial organization relative to β2 integrin activation conformation during spreading. (A)** Representative images of talin-1 (EGFP) and β2 integrin high-affinity conformation (mAb24) during the spreading phase. **(B)** Representative images of talin-1 (EGFP) and extended β2 integrin conformation (KIM127) during the spreading phase. Cell boundaries are outlined.

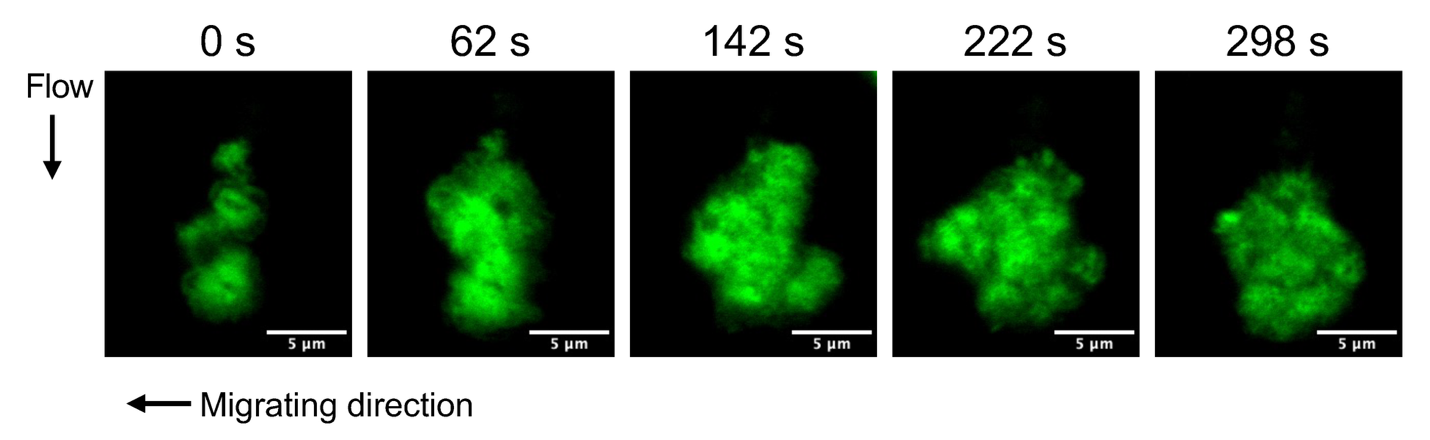

**Supplementary Figure 6.** **Additional representative neutrophil showing talin-1 redistribution during migration.** Representative time-lapse TIRF images of EGFP-talin1 during neutrophil migration under flow. Selected timepoints at the indicated are shown. Migration direction is indicated. Scale bar, 5 μm.

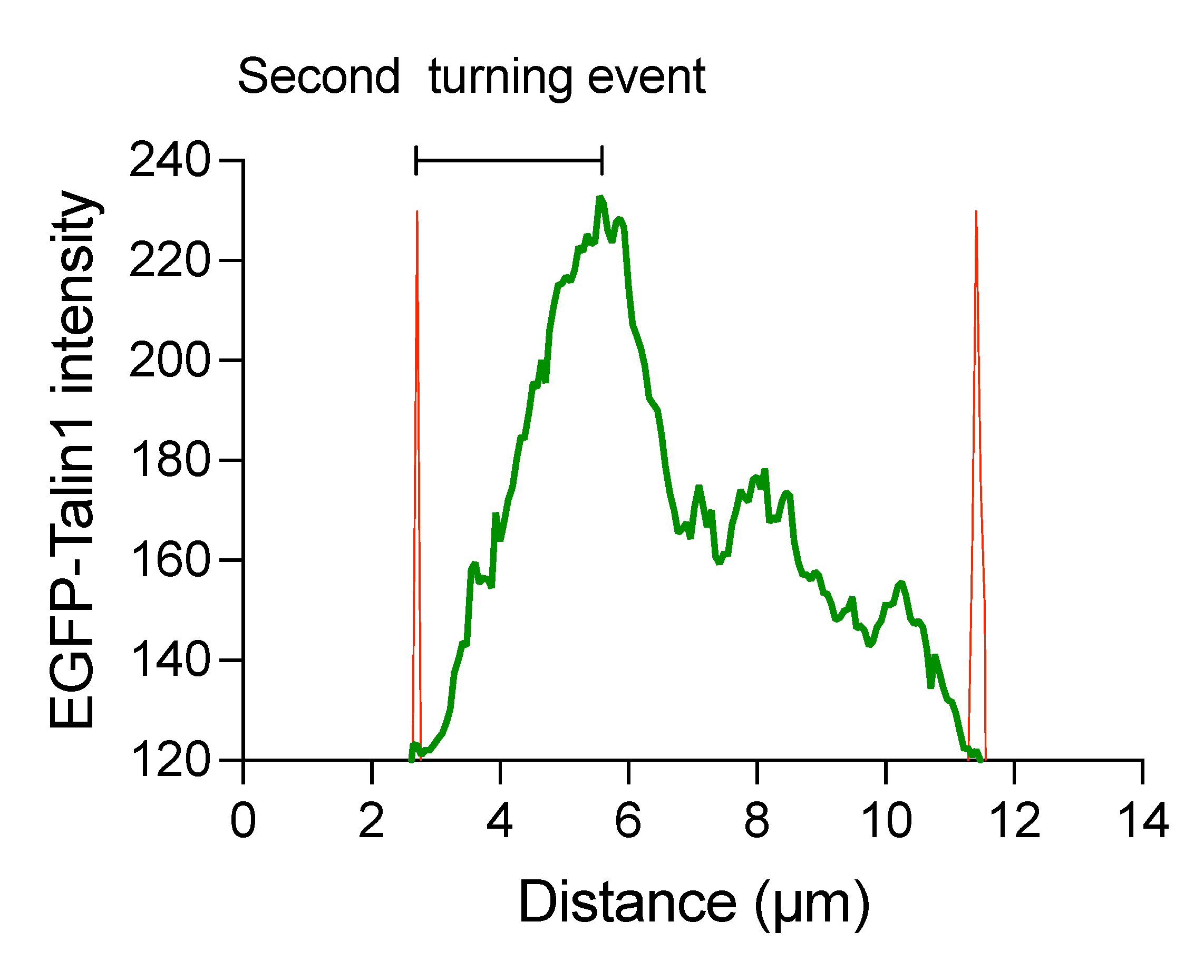

**Supplementary Figure 7. Line-scan analysis of a second turning event in the same cell.** A representative frame is shown. Consistent with Figure 4C, talin-1 enrichment precedes membrane extension toward the turning direction and is followed by spatial separation from the advancing cell edge. Distances between the talin-1 peak and the nearest cell edge were 2.52, 2.84, and 3.35 µm.

**Supplementary Video legends**

**Supplementary Video 1.** **Talin-1 accumulation and β2 integrin activation during neutrophil rolling and arrest.** Time-lapse TIRF imaging corresponding to Fig. 3A showing Talin-1 (EGFP) and β2 integrin activation (mAb24) during neutrophil rolling, arrest, and initial spreading. Bone marrow neutrophils from humanized β2 integrin (hITGB2) mice crossed with EGFP-talin1 knock-in mice were labelled with mAb24-DL550 and perfused over P-selectin and ICAM-1 coated surfaces under shear flow in the presence of CXCL1. Talin-1 (left), mAb24 (middle), and merged channels (right) are shown over time. Flow direction is indicated. Time is shown in seconds. Scale bar, 5 μm.

**Supplementary Video 2.** **Talin-1 spatial organization relative to β2 integrin activation conformations during neutrophil spreading.** Time-lapse imaging corresponding to Fig. 3 C and D showing the spatial relationship of talin-1 with β2 integrin high-affinity conformation (mAb24; H+, top) and extended integrin (KIM127; E+, bottom) during the spreading phase. Cell boundaries are outlined. Talin-1 (left), mAb24 or KIM127 (middle), and merged channels (right) are shown over time. Time is shown in seconds.

**Supplementary Video 3.** **Talin-1 redistribution in a migrating neutrophil.** Time-lapse TIRF imaging of EGFP-talin1 in a neutrophil migrating under flow. The cell corresponds to the representative example shown in Fig. 4A. Talin-1 signal redistributes over time, forming dynamic accumulation sites during migration. Flow direction is indicated. Scale bar, 5 μm.

**Supplementary Video 4.** **A neutrophil completing transendothelial migration and uropod detachment.** This video corresponds to Fig. 5C. Intravital imaging of EGFP-talin1 (green) and Ly6G (red) showing a neutrophil completing transendothelial migration and fully detaching from the endothelium. Talin-1 redistributes dynamically during diapedesis and detachment. Time stamp in minutes : seconds. Scale bar, 20 μm.

**Supplementary Video 5.** **A neutrophil undergoing interstitial migration following extravasation.** An intravital video of EGFP-talin1 showing a neutrophil migrating within the interstitial tissue following extravasation. Time stamp in minutes : seconds. Scale bar, 10 μm.
